# The diverse diet of southern Alaska resident killer whales changes across spatiotemporally distinct foraging aggregations

**DOI:** 10.1101/2024.09.12.612612

**Authors:** Hannah Myers, Daniel Olsen, Amy Van Cise, Kim Parsons, Abigail Wells, Craig Matkin

## Abstract

Top predators influence ecological communities in part through the prey they consume, which they often track through cycles of seasonal and geographic abundance. Killer whales are top predators in the marine ecosystem. In the North Pacific, they have diverged into three distinct lineages with different diets, of which the fish-eating type is most abundant. In this study, we examine the diet of the southern Alaska resident killer whale population across three major foraging aggregations. We take advantage of two unique sampling methods to reveal strong spatiotemporal patterns in diet from May through September. Chinook, chum, and coho salmon were each dominant in different locations and times, with substantial dietary contributions from Pacific halibut, arrowtooth flounder, and sablefish. The diverse, location-specific, and seasonal nature of the feeding habits of this marine top predator highlights the importance of diet sampling across broad spatiotemporal and population-level scales.

## Introduction

Top predators influence ecological communities in part by affecting the abundance and behavior of prey species they consume. Many predators track prey resources according to their abundance, timing, ephemerality, and predictability, among other factors^1^. Both marine and terrestrial predators show pronounced seasonal patterns in foraging strategies, and individuals and family or social groups often further specialize^2–5^. Understanding changes in diet across time, space, and among conspecifics is therefore critical to assess the ecological effects of predators across ecosystems. Killer whales (*Orcinus orca*) are top predators in the marine ecosystem and are most abundant in high-latitude regions^6^. In the North Pacific, they have diverged into three distinct ecotypes that may represent separate species^7^: those that eat exclusively fish (known as residents), those that eat exclusively mammals (called transients or Bigg’s), and those that likely consume mostly sharks (known as offshores)^7,8^. The fish-eating type is most abundant, with at least four studied parapatric populations spanning coastal regions in the northeast Pacific^8^. The 75 animals in the critically endangered southern resident population are found primarily in the Salish Sea and off the outer Washington coast^9^. The northern resident population of more than 300 individuals is found primarily off the coasts of British Columbia and southeast Alaska^10^. The southern Alaska resident population of about 1,000 killer whales ranges from southeast Alaska to Kodiak Island^11^. Finally, roughly 1,000 animals in the western Alaska North Pacific population range from Kodiak Island into the Bering Sea^12^.

Pacific salmon (*Oncorhynchus spp*.) are important prey for all studied North Pacific fish-eating killer whale populations. Southern residents, in particular, feed almost exclusively on Chinook salmon (*Oncorhynchus tshawytscha*) in spring and summer—though their diet is significantly more diverse in fall and winter^13^. Northern residents forage mostly for Chinook salmon and chum salmon (*Oncorhynchus keta*) ^14^ in summer. Southern Alaska residents eat substantial portions of Chinook, chum, and coho salmon (*Oncorhynchus kisutch*)^15,16^. Western Alaska North Pacific residents are less well-studied, but may consume lower trophic level fish further west where salmon are less available^17^, and at least some pods depredate on groundfish species^18^. On the other side of the Pacific, fish-eating killer whales in Avacha Gulf eat mostly coho and chum salmon in summer^19^.

While the importance of salmon as prey for North Pacific fish-eating killer whales is well-estableshed, how these animals utilize different prey resources across space and time is largely undocumented—especially for the substantially larger populations found off Alaska. In this study, we take advantage of a long-term killer whale monitoring program to examine how the diet of southern Alaska resident killer whales changes across three main foraging aggregations. We use both prey (scale and flesh) samples and fecal samples to identify the relative importance of different prey species. This work adds to our understanding of prey selection by fish-eating killer whales, providing valuable insights into their spatiotemporal foraging strategies and the potential ecological impacts of prey variability.

## Results

We collected 255 samples of fish scales or flesh while southern Alaska resident killer whales were observed feeding across 31 years (1991 to 2021). We collected 186 fecal samples across six years (2016 – 2021), of which 87 successfully sequenced, were not from the same individual on the same day, and passed the quality check. These samples were collected in three adjacent geographic areas with largely non-overlapping data collection periods: Kenai Fjords from mid-May to mid-June, eastern Prince William Sound from mid-June to July, and western Prince William Sound from July through September (Figure 1). Both prey (scale and flesh) and fecal samples demonstrated distinct dietary patterns across these three foraging aggregations. Chinook and chum salmon were the dominant species across all diet samples, although coho salmon were the most common prey sample collected in western Prince William Sound. Fecal samples also revealed substantial contributions from Pacific halibut (*Hippoglossus stenolepis*), arrowtooth flounder (*Atheresthes stomias*), and sablefish (*Anoplopoma fimbria*).

**Figure 1.**
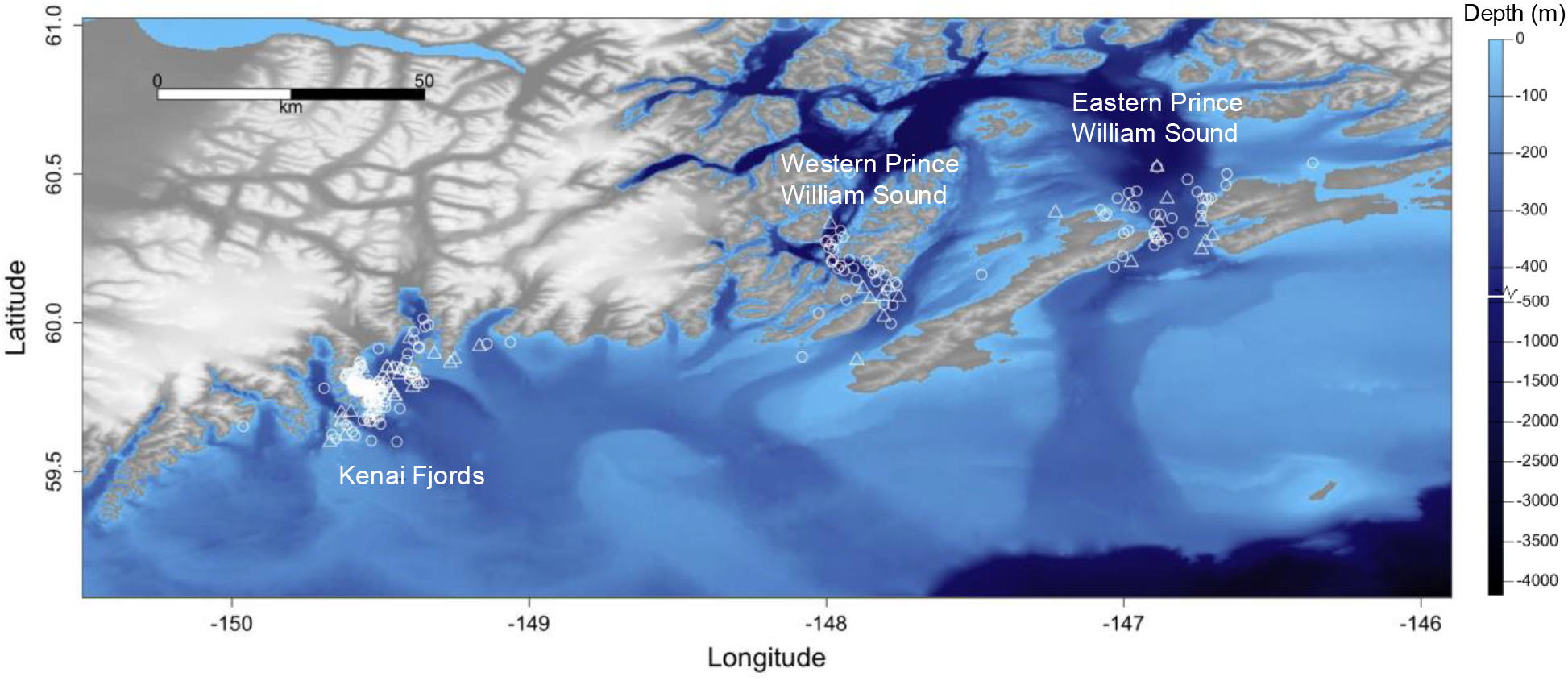
Map of prey (circle) and fecal (triangle) samples collected in the study area in the northern Gulf of Alaska.

### Prey samples

Of the 255 scale and flesh samples we collected, 249 were of three salmon species: Chinook, chum, and coho (Table 1, Figure 2). Only one prey sample did not clearly align with an aggregation area and was therefore removed from the analysis (chum salmon collected September 4^th^, 2020 at 59.788° N, 148.705° W). Chinook salmon made up 77% of prey samples collected in Kenai Fjords—where 80% of samples were collected between May 17^th^ and June 12^th^. Chum salmon were the primary prey species in eastern Prince William Sound (62% of samples), where sampling took place primarily from June 15^th^ to July 22^nd^. Coho salmon dominated the western Prince William Sound foraging aggregation (77%), where we collected samples primarily from July 22^nd^ to September 10^th^. In addition to the three main salmon prey species, five sockeye salmon (*Oncorhynchus nerka*) prey samples were collected in May and June in Kenai Fjords and eastern Prince William Sound, and one Pacific herring (*Clupea pallasii*) sample was collected in early May in Kenai Fjords. The main pods (maternally related family groups) differed across the three foraging aggregations, although samples were collected from the AK pod across all three (Table 1).

**Table 1.** Fish scale and flesh samples from three primary salmon prey species, collected while killer whales were observed feeding. The period within which 80% of prey samples were collected and the range of dates within which all prey samples were collected are shown for each area. The main pods are those from which at least 5% of samples were collected, with the number of samples from that pod in parentheses. The predominant prey species from each foraging aggregation is shown in bold.

| Area | Total samples | Sampling period (80% of samples / all samples) | Main pods (*n*) | Salmon species | Proportion of samples |
| --- | --- | --- | --- | --- | --- |
| Kenai Fjords | 149 | May 17 – Jun 12 / May 3 – Sep 8 | AD8 (41), AK (37), AD5 (24), AD16 (16) | **Chinook** | **.78** |
|  |  |  |  | Chum | .19 |
|  |  |  |  | Coho | .03 |
| Eastern Prince William Sound | 37 | Jun 15 – Jul 21 / Jun 14 – Aug 29 | AB (7), AD8 (4), AE (4), AJ (4), AK (4), AI (3), AX48 (2) | Chinook | .16 |
|  |  |  |  | **Chum** | **.62** |
|  |  |  |  | Coho | .22 |
| Western Prince William Sound | 62 | Jul 22 – Sep 9 / Apr 29 – Sep 21 | AE (26), AB (9), AK (8), AI (4) | Chinook | .19 |
|  |  |  |  | Chum | .03 |
|  |  |  |  | **Coho** | **.77** |

**Figure 2.**
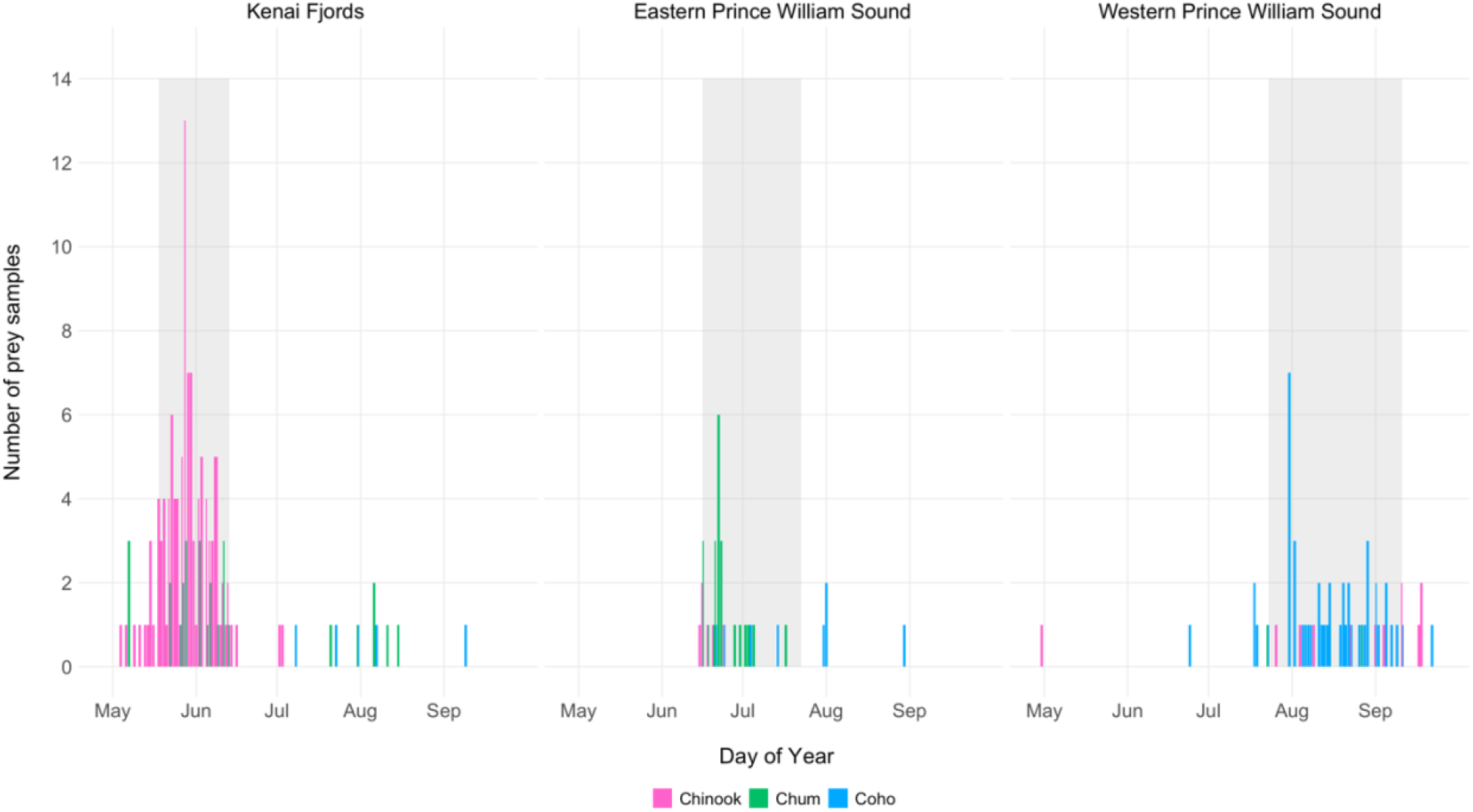
Prey samples collected from each area from May through September across all years. Gray shaded periods indicate the days in which 80% of prey samples were collected in each area.

We used a Bayesian multinomial logistic regression model to test for relationships between prey species and foraging aggregation. Model results reflected the strong diet patterns described above (Table 2). Coefficients for which the 95% credible interval does not include zero may be interpreted as statistically significant. In particular, the strong quadratic effect for chum salmon reflects the high probability of chum salmon in eastern Prince William Sound, and the strong linear effect for coho salmon reflects the higher probability of coho salmon in western Prince William Sound. Posterior predictive checks indicated good model fit to the data and model convergence and 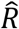 was 1.00 for all parameters.

**Table 2.** Prey sample multinomial logistic regression model results for the probability of Chinook, chum, and coho salmon prey as a function of aggregation area. The model reference level is Chinook salmon. The linear term describes the relationship across the three ordered areas: Kenai Fjords, eastern Prince William Sound, and western Prince William Sound. The quadratic term reflects how the middle level (eastern Prince William Sound) differs from the expected linear relationship between the first (Kenai Fjords) and last (western Prince William Sound) levels. Statistically significant effects (those for which the 95% credible interval does not include zero) are shown in bold.

| Species | Coefficient | Estimate | Estimated error | Lower 95% credible interval | Upper 95% credible interval |
| --- | --- | --- | --- | --- | --- |
| Chum salmon | Intercept | **-0.66** | **0.33** | **-1.36** | **-0.06** |
|  | Linear | -0.39 | 0.60 | -1.72 | 0.65 |
|  | Quadratic | **-2.55** | **0.53** | **-3.66** | **-1.59** |
| Coho salmon | Intercept | -0.49 | 0.27 | -1.03 | 0.04 |
|  | Linear | **3.28** | **0.40** | **2.55** | **4.12** |
|  | Quadratic | -0.99 | 0.51 | -2.01 | 0.01 |

### Fecal samples

Results from 87 fecal samples reinforced the primary importance of salmonids in the diet of southern Alaska resident killer whales while also revealing three major prey items that are presumably captured and consumed at depth: Pacific halibut, arrowtooth flounder, and sablefish (Table 3, Figure 3). As with prey samples, Chinook salmon was the dominant species in fecal samples from Kenai Fjords (71%) and chum salmon was the dominant species detected in eastern Prince William Sound (72%). In western Prince William Sound, diet samples were substantially more diverse, with the greatest contribution from Chinook salmon (35%). The main sampling periods largely aligned for both prey and fecal samples, however, fecal samples were collected in fewer years and sample sizes were notably lower, especially from Prince William Sound (Tables 1 and 3).

**Table 3.** Fecal samples from six main fish prey species. The period within which 80% of fecal samples were collected and the range of dates within which all fecal samples were collected are shown for each area. The species with the highest overall proportion from each foraging aggregation is shown in bold.

| Area | Total samples | Sampling period (80% of samples / all samples) | Pods identified (*n*) | Major prey species | Proportion across all samples |
| --- | --- | --- | --- | --- | --- |
| Kenai Fjords | 66 | May 24 – Jun 8 / May 17 – Sep 17 | AD8 (21), AK (18), AD16 (10), AJ (1), AX48 (1) | **Chinook** | **.71** |
|  |  |  |  | Chum | .26 |
|  |  |  |  | Coho | .00 |
|  |  |  |  | Halibut | .03 |
|  |  |  |  | Arrowtooth | .00 |
|  |  |  |  | Sablefish | .00 |
| Eastern Prince William Sound | 12 | Jun 15 – Jun 30 / May 9 – Jul 1 | AK (3), AE (1), AJ (1), AX48 (1) | Chinook | .04 |
|  |  |  |  | **Chum** | **.72** |
|  |  |  |  | Coho | .02 |
|  |  |  |  | Halibut | .07 |
|  |  |  |  | Arrowtooth | .13 |
|  |  |  |  | Sablefish | .03 |
| Western Prince William Sound | 9 | Jul 3 – Sep 30 / Jul 2 – Sep 30 | AK (2), AB25 (1), AE (1), AI (1), AJ (1) | **Chinook** | **.35** |
|  |  |  |  | Chum | .20 |
|  |  |  |  | Coho | .12 |
|  |  |  |  | Halibut | .24 |
|  |  |  |  | Arrowtooth | .09 |
|  |  |  |  | Sablefish | .00 |

**Figure 3.**
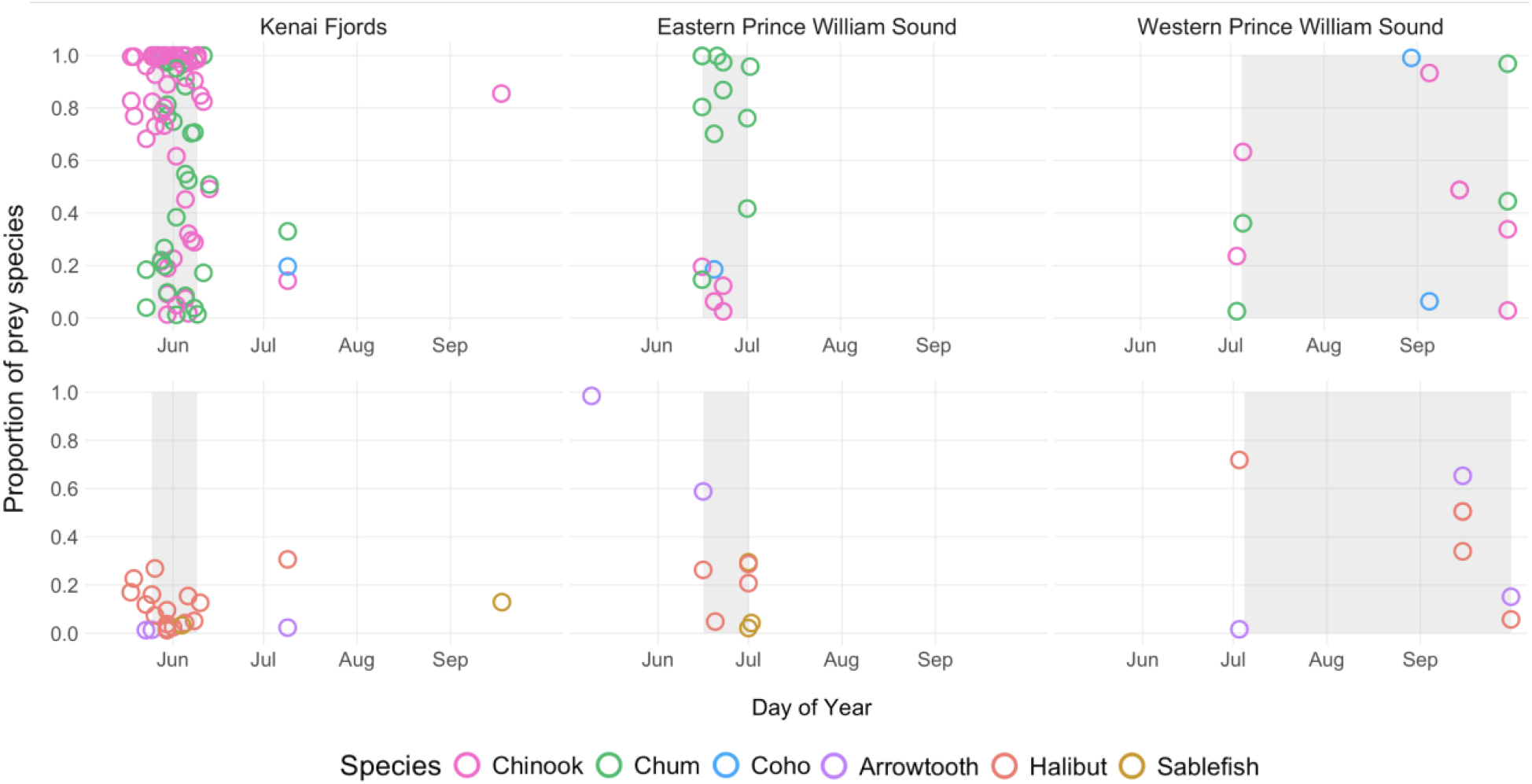
Proportion of prey species in fecal samples collected in each area from May through September. Chinook, chum, and coho salmon are in the top panel and Pacific halibut, arrowtooth flounder, and sablefish are in the bottom panel. Gray shaded periods indicate the days in which 80% of fecal samples were collected in each area.

Host DNA from 48 unique whales was identifiable in 82 samples and samples originating from the same host identified. Pod identity was assigned in 63 cases based on photo-identification at the time of sampling (Table 3). As with prey samples, fecal samples were collected from the AK pod across all three foraging aggregations, though there were differences in some of the other pods sampled across events.

Halibut was detected in all regions and made up >5% of at least one sample in May, June, July, and September. It was consumed regularly by the AE, AK, AD8, and AD16 pods.

Arrowtooth flounder was detected in the greatest proportions in May, June, and September, primarily in samples from the AE pod. In addition to the six major prey species, three fecal samples contained 1-3% prowfish (*Zaprora silenus*), all of which were from Kenai Fjords.

For fecal samples, we fit a Bayesian multinomial model with a Dirichlet response distribution to describe the effect of foraging aggregation on the probabilities of the six main prey species (Table 4). The strongest effects were evident for chum and Chinook salmon: the high proportion of chum salmon in eastern Prince William Sound (represented by the intercept) and the high proportion of Chinook salmon in Kenai Fjords were statistically significant (i.e., the 95% credible intervals did not include zero). Flatfish were more likely in western Prince William Sound, though not significantly so. Other species’ proportions varied but effects were not consistently significant. Posterior predictive checks indicated sufficient model fit to the data and model convergence and 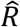 was 1.00 for all parameters.

**Table 4.** Fecal sample model results for the probability of six main prey species as a function of foraging aggregation. Sablefish is the reference species and eastern Prince William Sound is the reference area against which intercepts and coefficients for Kenai Fjords (KF) and western Prince William Sound (WPWS) are compared. Statistically significant effects (those for which the 95% credible interval does not include zero) are shown in bold.

| Species | Coefficient | Estimate | Estimated error | Lower 95% credible interval | Upper 95% credible interval |
| --- | --- | --- | --- | --- | --- |
| Arrowtooth flounder | Intercept | -0.15 | 0.42 | -1.00 | 0.65 |
|  | KF | 0.29 | 0.45 | -0.61 | 1.18 |
|  | WPWS | 0.77 | 0.64 | -0.52 | 2.01 |
| Pacific halibut | Intercept | 0.28 | 0.41 | -0.55 | 1.09 |
|  | KF | 0.21 | 0.45 | -0.67 | 1.09 |
|  | WPWS | 0.81 | 0.62 | -0.41 | 2.02 |
| Chum salmon | Intercept | **1.95** | **0.35** | **1.28** | **2.67** |
|  | KF | **-1.20** | **0.38** | **-1.97** | **-0.46** |
|  | WPWS | **-1.34** | **0.58** | **-2.49** | **-0.25** |
| Coho salmon | Intercept | -0.20 | 0.41 | -1.02 | 0.61 |
|  | KF | 0.17 | 0.44 | -0.71 | 1.04 |
|  | WPWS | 0.44 | 0.63 | -0.80 | 1.67 |
| Chinook salmon | Intercept | 0.47 | 0.42 | -0.34 | 1.31 |
|  | KF | **1.76** | **0.44** | **0.88** | **2.64** |
|  | WPWS | 1.17 | 0.60 | -0.01 | 2.35 |

#### Additional observations

The rich data available from fecal samples revealed several unique observations that complement the data generated from prey scales and tissue remains. First, fecal samples were collected across multiple foraging aggregations from four individual whales, all from the AK pod. Consistent with overall results, samples from these individuals included mostly Chinook salmon during the Kenai Fjords foraging aggregation and mostly chum salmon during the eastern Prince William Sound aggregation. Second, on May 29^th^, 2018, fecal samples were collected from both adult female AD31 and her 4-year-old male offspring AD54, in the AD8 pod. Samples from both mother and offspring contained the same four species in very similar percentages: 81% and 77% chum salmon, respectively; 9% and 19% Chinook salmon, 9% and 4% Pacific halibut, and <1% each arrowtooth flounder. This unique collection event highlights an instance of high diet similarity between a known mother and dependent offspring—a pair of animals likely to share prey^20^. Third, in 2018, we observed a change in foraging patterns with killer whales foraging farther offshore than typical in Kenai Fjords. We documented a notably higher proportion of chum salmon (76%) in the 14 fecal samples collected in Kenai Fjords that year.

## Discussion

In this study, we found that southern Alaska resident killer whales utilized different prey resources across spatiotemporally distinct foraging aggregations. The shift from primarily Chinook to chum to coho salmon across these aggregations likely reflects the relative availability of these three priority prey resources. We also found that, even from May to September when salmon are abundant, southern Alaska resident killer whales supplement their diet with other high-energy-content fishes—particularly Pacific halibut, arrowtooth flounder, and sablefish.

In general, the strong agreement between prey and fecal samples in terms of primary species, timing, and pods involved reinforced the consistency of these killer whale foraging aggregations. A notable exception is the prevalence of coho salmon in prey samples from the western Prince William Sound foraging aggregation (77%) relative to its proportion in fecal samples from that area (12%). The low number of fecal samples from western Prince William Sound (*n* = 9) makes interpretation challenging; however, most prey samples were collected in late July and early August (Figure 2), whereas no fecal samples were collected during that time (Figure 3). In addition, 80% of prey samples from western Prince William Sound were collected prior to 2008, while all fecal samples were collected after 2016, so changes in prey availability or preferences over time may also be relevant. While interannual differences in killer whale diet were not specifically investigated as part of this study, the disproportionately high level of chum salmon detected in Kenai Fjords in 2018 suggests that they are likely. Although few fecal samples were collected in eastern Prince William Sound, the species composition was closely aligned for fecal (*n* = 12) and prey (*n* = 37) samples from that foraging aggregation.

Pacific salmon are an abundant yet locally ephemeral resource in this region. Tracking the spatiotemporal variation in their phenology—including across species—extends foraging opportunities on this important prey resource for consumers. For example, the distribution of coastal brown bears (*Urus* arctos) and glaucous-winged gulls (*Larus glaucescens*) reflects the shifting distribution of spawning sockeye salmon (*Oncorhynchus nerka*) within a watershed^21^. Individual brown bears visit multiple salmon spawning sites in synchrony with their spawning phenology^22^—and those that track spawning phenology for the longest consume the most salmon^4^. Though the pulsed nature of salmon availability in the marine environment is less well-defined, the results of this study suggest that fish-eating killer whales may also capitalize on changes in Pacific salmon availability across relatively short spatial and temporal scales.

Populations with diverse diets may include specialized individuals or groups that feed on a restricted subset of prey^23^. The distinct prey patterns we documented may reflect killer whales tracking prey resources as well as different prey preferences among pods, which may be culturally transmitted^24^. The main pods detected in this study generally have different core use areas^25^ and differ genetically^26^. The Kenai Fjords aggregation was dominated by pods that have a maternal haplotype shared with southern resident killer whales, while the pods encountered in eastern Prince William Sound mostly have a haplotype shared with the northern residents, and the pods commonly encountered in western Prince William Sound are more mixed. We found much higher proportions of flatfish in samples from the AE pod, which may indicate a slight degree of specialization for that pod. Almost all samples that included flatfish were from pods with the southern resident haplotype, though few fecal samples for which pod was identified were of the northern resident haplotype (*n* = 4). In contrast, the fecal samples from the AK pod matched the overall trend of high proportions of Chinook salmon during the Kenai Fjords aggregation and high chum salmon during the eastern Prince William Sound aggregation. On a finer scale, foraging strategies differ among sex- and reproductive classes of southern and northern resident killer whales^27^. While not a focus of this study, these classes are worthy of investigation.

The shifts in diet we documented for southern Alaska resident killer whales occurred within a limited season (May to September). However, seasonal changes in diet throughout the year are also common among top predators—even those with highly specialized foraging strategies. For example, wolves (*Canis lupus*) on Yellowstone National Park’s northern range whose diets are dominated by elk (*Cervus elaphus*) hunt more mule deer (*Odocoileus hemionus*) and bison calves (*Bison* bison) in spring and summer. In Alaska, some wolves that hunt ungulates also eat large portions of Pacific salmon in summer^3^.

Migrating gray whale (*Eschrictius* robustus) calves traveling through a geographic pinch point into the Bering Sea provide a pulse of high-quality prey that attracts mammal-eating killer whales in spring^2^. These killer whales change to a diet of largely northern fur seals (*Callorhinus ursinus*) in summer^28^. As previously mentioned, southern resident killer whales eat almost exclusively Chinook salmon in spring and summer, but in fall and winter up to about half of their diet is made up of chum and coho salmon, steelhead (*Oncorhynchus mykiss*), lingcod (*Ophiodon elongatus*), big skate (*Rana binoculata*), and flatfishes^13^.

In the northeast Pacific, fish-eating killer whale diet sampling has been highly biased to summer months and shallow coastal regions where salmon are likely to be abundant— including in this study^14,16,29^. At present, southern Alaska residents’ primary prey resources for more than half the year (October through April) remain unknown. Our current research effort likely misses other major foraging aggregations, even during the field season. The southern Alaska resident population is acoustically detected most often and with the largest estimated group sizes in late fall and winter in western Prince William Sound, and the peak in use of eastern Prince William Sound begins in early spring^30,31^—periods when no diet sampling took place. Lower quality or more difficult to access prey (such as flatfish) may be disproportionately important to killer whales if the their phenological patterns mean that they are available when higher quality prey (such as Chinook salmon) are not^1^.

However, abundant but notably smaller and lower calorie fishes—especially pink salmon (*Oncorhynchus gorbuscha*) and herring—were not detected in proportions >1% in any fecal sample in this study. It is possible that the small proportion of prowfish found in fecal samples and the single herring prey sample we collected were secondary prey (i.e., a fish eaten by killer whale prey). Prowfish are a documented prey item of Pacific halibut, but not of Pacific salmon, arrowtooth flounder, or sablefish^32,33^.

Similarly, pooling diet samples without accounting for spatiotemporal differences in sampling effort may mask important patterns^34^. For example, in this study, 59% of prey samples and 76% of fecal samples were collected in Kenai Fjords. Although sampling in Kenai Fjords took place for the shortest period of time, the protected waters in this area created better conditions to search for samples. The smaller killer whale group sizes encountered in this location enabled faster photo-identification, allowing more search time for diet samples. Our diet data is therefore overweighted with samples that contain a high proportion of Chinook salmon from this relatively short foraging aggregation, and if samples were not separated by aggregation the importance of chum and coho salmon at other places and times would be diluted. Additionally, collecting fish scales or pieces of flesh when killer whales are observed feeding at the surface biases results toward salmonids, while species that are captured and consumed at depth (and those without scales) go undetected. DNA analysis of fecal samples has revealed a greater diversity of fish species consumed by resident killer whales^13,15,35^.

Assumptions about the diets of predator populations are frequently drawn from data collected over brief periods in specific regions, often focusing on unique (sub)populations that may not represent broader spatiotemporal or species-level scales. Samples collected in summer in the Salish Sea from southern resident killer whales are overrepresented in North Pacific fish-eating killer whale diet studies^36–38^. Extrapolating the diet of this endangered population across three populations several orders of magnitude more abundant may lead to problematic conclusions with potential management implications^36–38^. This study adds a robust analysis to the growing body of diet research from other fish-eating killer whale populations in the North Pacific^15,19,39^.

Finally, predator populations with highly specialized diets are likely to be more vulnerable to disturbance—including climate change impacts—than more generalist predators with greater flexibility in their diets^40,41^. We found that southern Alaska resident killer whales utilized three different primary salmonid prey resources across three main summer foraging aggregations, with substantial supplementation from other fishes. This suggests that the southern Alaska resident population—which is thought to be growing at a rate near maximum^11^—has a more diverse summer diet than that of the endangered southern resident population^13^. Specializing on very high trophic level fish (e.g., Chinook salmon) is also likely to limit population growth compared to eating a more diverse diet that includes relatively lower trophic level fishes. However, whether the diet of southern Alaska resident killer whales remains more diverse year-round has yet to be studied.

## Methods

### Sample collection

Fieldwork took place between May and September in Prince William Sound and Kenai Fjords, Alaska (Figure 1). Prey samples were collected from 1991 to 2021 and fecal samples were collected from 2016 to 2021. Prince William Sound is characterized by large barrier islands, entrances with strong tidal currents, and small islands scattered near glacially carved trenches. Kenai Fjords runs along the southern Alaska coast and has many long glacially carved fjords. All fieldwork was conducted from a 11 m research vessel concurrent with other research tasks including photo-identification, acoustic research, and body condition assessment.

Prey samples from predation events and fecal samples were collected from surface waters during focal follows of resident killer whale groups. Predation events were often identified when killer whales made tight turns at the surface and/or when a whale approached to share fish with its mother. Scales or tissue samples were typically found in upwellings (upward currents from the flukes of a diving whale) after observing that a fish had been shared or broken apart at the surface. Fecal samples were usually found in upwellings while the vessel was travelling >100 m behind the whales. Samples were collected from the bow using a long-handled fine-mesh dip net and transferred into new glass jars. All samples were labelled and frozen within 10 min of collection; prey samples were frozen in ethanol.

The individuals, pods, and number of killer whales involved in each encounter in which a diet sample was collected was determined using photo-identification. Identification photos were taken with a Nikon D700 or D750 camera with a 300 or 400 mm lens and matched to a long-term photo-identification catalog maintained by the North Gulf Oceanic Society^42^. In each encounter, all members of a matriline (defined as a mother or grandmother and her offspring) were assumed to be present if at least one member of the matriline was photographed, as in Olsen *et al*.^43^ and Myers *et al*.^44^.

### Prey species identification

Prey samples were identified to species using fish scale morphology or genetic analysis, as in Withler *et al*.^45^, Ford and Ellis^14^, Hanson *et al*.^29^. Only one sample per hour was retained for analysis to avoid pseudoreplication. For fecal samples, prey species composition and host identity were determined using the genetic techniques described below.

#### Fecal sample sequencing

Whole genomic DNA was extracted from a pea-sized subsample of frozen fecal matter using the QIAamp^®^ Fast DNA Stool mini kit following standard protocols executed using the QIAcube automated extraction robot^35^. 16S SSU rDNA was targeted using custom-designed Illumina primers for salmon and groundfish, as previously published for the prey metabarcoding of southern resident killer whales^35^. Amplification reactions contained 4uL of DNA, 1X Promega GoTaq Flexi buffer (Promega Corp., Madison WI), 3.0mM MgCl2, 0.2mM of each dNTP, 0.1ug/uL of BSA, 0.2uM of each primer, and 2 units of Promega GoTaq Flexi DNA Polymerase. Communities were amplified in a 32-cycle PCR, with cycling conditions as follows: initial denaturation at 94°C for 2 min, followed by 32 cycles of 94°C for 35 sec; 61°C for 1 min; 72°C for 35 sec; and a final extension at 72°C for 5 min. Amplicons were gel cleaned using Qiagen MinElute columns to remove non-target PCR products and primer dimer.

Cleaned amplicons were individually indexed using two different sets of indices. In 2018 and 2019, Illumina Nextera forward and reverse index tags were used, which create a unique combination of indices by using a unique forward primer for each column and a unique reverse primer for each row on a 96-well plate. This indexing PCR was completed using a 50uL reaction containing 8uL of gel purified PCR product, 1X NEB Phusion High-Fidelity master mix (New England BioLabs), 0.2mM of each dNTP, 5uL each of one Illumina Nextera forward and reverse index tag. In the two 2021 MiSeq runs, Illumina Nextera DNA Unique Dual Index (UDI) primers were used (Illumina, Inc.), comprising unique forward and reverse indexes for each well in a 96-well plate in order to reduce the effect of index hopping^46^. The index PCR was performed in a 40ul reaction containing 8uL of gel purified PCR product, 1uL each of one Illumina Nextera UDI forward and reverse index, and 30uL of Phusion High-Fidelity PCR Master Mix (New England Biolabs). Regardless of the indices used, the indexing PCR conditions remained the same: 72°C for 3 min, 98°C for 30 sec, followed by 12 cycles of 98°C for 10 sec, 55°C for 30 sec, 72°C for 30 sec, and a final extension at 72°C for 5 min. Samples were sequenced on four sequencing runs (2018, 2019, and two in 2021) using an Illumina MiSeq next generation sequencer (Illumina, Inc.) at the Northwest Fisheries Science Center, NOAA Fisheries, Seattle, WA.

In addition to fecal samples from southern Alaska resident killer whales, two mock communities including pre-determined quantities of genomic DNA from several vouchered fish species were included on each of the four sequencing runs to detect and control for any potential bias caused by index hopping, species-specific amplification efficiency, or genotyping error. Details on mock community generation can be found in Van Cise *et al*.^15^. Mock communities were sequenced alongside sample libraries each year.

#### Fecal sample sequence alignment and QAQC

Sequences from all runs were combined and analyzed using a custom pipeline based on the dada2 package^47^ in the R computing environment^48^. This pipeline includes steps for (1) trimming sequences based on general sequence quality, (2) filtering sequences based on a maximum number of expected errors^49^, (3) learning the error rates for each possible transition, (4) de-replicating sequences by combining and counting identical sequencing reads to reduce computation time, and finally inferring unique amplicon sequence variants (ASVs) from the filtered and trimmed sequences using the previously learned error rates^47^. Once unique ASVs were identified, paired forward and reverse reads were merged and chimeras removed. Taxonomy was assigned to the remaining ASVs using a naïve Bayesian classifier^50^ that relies on a fasta-formatted reference database, which we custom-generated by downloading sequences for all fish and shark species from NCBI GenBank.

Because various sources of laboratory-introduced bias can affect the observed number of reads assigned to a given species, mock community control samples were used to estimate and correct for the effects of errors, e.g. from amplification bias and index hopping. This model was run using both mock communities to estimate species-specific bias and read proportions by species were corrected based on model estimates (Figure 2, Van Cise *et al*.^15^). Some prey species in the final dataset were not anticipated, e.g. sablefish, and therefore not included in the mock communities and could not be corrected. However, overall differences between uncorrected and corrected proportional data are small enough that the difference for these species is expected to be minor (Supplemental Figures S2 and S3, Van Cise *et al*.^15^).

Final data filtering consisted of removing ASVs that assigned to *Orcinus orca* and aggregating ASVs by species. Laboratory and field duplicates used to track potential sources of bias or contamination were removed from the dataset before analysis. Samples with a read depth < 25,000 reads were removed from the dataset. Additionally, individual whale ID was genetically determined using a previously developed panel of SNPs (see Van Cise *et al*^15^. for methodological details), and samples were removed if they were collected from the same individual on the same day to avoid pseudoreplication. Prey species were only included in downstream analyses if they represented >1% of the reads in one or more samples in the dataset to avoid potential bias from genotyping error.

## Statistical analyses

We modeled the effect of foraging aggregation on prey species composition with Bayesian multinomial regression models. For prey samples we used a multinomial logistic regression, for fecal samples we used a Dirichlet-multinomial distribution to handle the proportional nature of the data. We used the area in which the sample was collected (Kenai Fjords, eastern Prince William Sound, or western Prince William Sound) as a three-factor categorical variable to represent the foraging aggregation. Models were implemented using the *brms* package^51^ in R (version 4.3.3) with the default uninformative priors.

## Acknowledgements

Graeme Ellis completed photo-identification analysis. Fish prey species identification was completed by the Sclerochronology Laboratory and Molecular Genetics Laboratory at the Pacific Biological Station, Department of Fisheries and Oceans, Nanaimo, B.C., Canada, with special thanks to Brianna Wright. Funding for sample analysis was provided by the National Fish and Wildlife Federation through a research grant awarded to K.M. Parsons and the North Gulf Oceanic Society. Funding to support sample collection was provided by the *Exxon Valdez* Oil Spill Trustee Council and the U.S. Marine Mammal Commission.

## Notes

### Competing Interest Statement

The authors have declared no competing interest.

